# Y-box binding protein 1 interacts with dengue virus nucleocapsid and mediates viral assembly

**DOI:** 10.1101/2022.01.25.477802

**Authors:** Mayra Diosa-Toro, Debbie R Kennedy, Vanessa Chuo, Vsevolod L. Popov, Julien Pompon, Mariano A. Garcia-Blanco

**Author notes:** Department of Biomolecular Health Sciences, Faculty of Veterinary Medicine, Utrecht University, The Netherlands (Mayra Diosa-Toro). Address correspondence to Mariano A. Garcia-Blanco.

## Abstract

Infection with dengue virus (DENV) induces vast rearrangements of the endoplasmic reticulum, which allows the compartmentalization of viral RNA replication and particle assembly. Both processes occur in concert with viral and cellular proteins. Prior studies from our group suggest that the host RNA-binding protein (RBP) Y-box binding protein 1 (YBX1) is required for a late step in the DENV replication cycle. Here we report that YBX1 interacts with the viral nucleocapsid, distributes to DENV assembly sites and is required for efficient assembly of intracellular infectious virions and their secretion. Genetic ablation of YBX1 decreased the spatial proximity between capsid and envelope, increased the susceptibility of envelope to proteinase-K mediated degradation, resulted in the formation of rough empty-looking particles, and decreased the secretion of viral particles. We propose a model wherein YBX1 enables the interaction between the viral nucleocapsid with the structural protein E, which is required for proper assembly of intracellular virus particles and their secretion.

**Importance:** The global incidence of dengue virus (DENV) infections has steadily increased over the past decades representing an enormous challenge for public health. During infection, DENV viral RNA interacts with numerous host RNA binding proteins (RBPs) that aid viral replication and thus constitute potential molecular targets to curb infection. We recently reported that Y-box-binding protein 1 (YBX1) interacts with DENV RNA and is required at a late step of the replication cycle. Here we describe the molecular mechanism by which YBX1 mediates DENV infection. We show that YBX1 interacts with the viral nucleocapsid, distributes to DENV assembly sites and is required for efficient assembly of intracellular infectious virions. These results provide important insights into DENV assembly, revealing novel functions of host RBPs during viral infection and opening new avenues for antiviral intervention.

## INTRODUCTION

Despite massive efforts in the fight against dengue, the global incidence of this arthropod-borne viral disease has grown dramatically in recent decades. The etiological agent, the four dengue viruses (DENV1-4), are endemic to more than 100 countries, where annual outbreaks result in high disease and economic burden (1). There are no specific antiviral therapies available against dengue and, although the only licensed vaccine reduces the incidence of severe disease, serious concerns regarding its overall efficacy and safety remain (2).

DENV belongs to the *Flavivirus* genus, which encompasses other arboviruses of clinical importance such as Zika and West Nile viruses. Flaviviruses carry a single stranded positive-sense RNA genome packaged by the viral protein capsid (C). This nucleocapsid is surrounded by a lipid bilayer derived from the host cell in which, the viral glycoproteins envelope (E) and membrane (M) are found. Upon delivery into a susceptible cell, the genomic viral RNA (vRNA) is translated into the viral proteins (3). The vRNA also functions as a template for replication via the synthesis of a complementary negative-sense strand and, thus, the formation of double stranded RNA (dsRNA) intermediates. Replication takes place in virus-induced endoplasmic reticulum (ER) invaginations, which are juxtaposed to but distinct from the assembly sites (4). Hence, single stranded RNA genomes must be relocated from the replication to the assembly sites. During assembly, virions, referred to as immature, bud into the ER lumen and then traffic to the *trans* Golgi apparatus where the cellular protease furin cleavage the pr peptide from M, thus generating mature virions (5). These reach the extracellular milieu via the secretory pathway. Throughout these processes DENV vRNA has been shown to interact with multiple host RNA-binding proteins (RBPs), which can either promote viral replication or are part of the antiviral arsenal of the host cell (recently reviewed in (3, 6, 7)). One of these RBPs is Y-box-binding protein 1 (YBX1 or YB-1), which was found to bind to DENV vRNA in cell free reactions and in DENV-infected cells (8, 9).

YBX1, YBX2 and YBX3 form the Y-box protein family, the predominant group of cold-shock domain (CSD) proteins present in humans (10). CSD proteins are highly conserved nucleic-binding proteins with pleiotropic functions in DNA and RNA-dependent processes (11–14). YBX1 is composed of three structural domains, the N-terminal alanine/proline-rich (A/P) domain, the CSD and the C-terminal domain (CTD) that contains positively and negatively charged clusters of amino acids (14). Binding to nucleic acids is mediated by two consensus motives, RNAP1 and RNAP2, localized in the CSD. YBX1 is found in the nucleus as well as in the cytoplasm. In the nucleus, binding of YBX1 to DNA significantly decreases the melting temperature of double helices enabling transcription and DNA repair (12). In addition, by binding to exonic enhancers, YBX1 participates in the splicing of several pre-mRNAs, and this has been implicated in the development and persistence of malignancies (15–17). In the cytoplasm, YBX1 is a highly abundant component of cytoplasmic messenger ribonucleoproteins (mRNPs) and is a key regulator of mRNA transport, translation and decay (12, 18, 19). YBX1 is also present in RNA-containing condensates such as stress granules and P-bodies (20, 21), and plays a key role in determining the RNA composition of extracellular vesicles by sorting short non-coding RNAs into exosomes (22–25).

Given its numerous functions during RNA metabolism, it is not surprising that by binding to vRNA genomes, YBX1 also regulates the outcome of virus infection. For example, YBX1 knockdown prior to transfection with an infectious clone of hepatitis C virus (HCV), caused an increase in the progeny virus, in spite of a decrease in intracellular infectivity (26, 27). Comparably, DENV-infected mouse embryonic fibroblasts (MEFs) derived from YBX1 knockout (KO) mice, were shown to produce higher virus titers than their wild-type (WT) counterparts (8). These studies suggest an antiviral function for YBX1; however, other studies point to a proviral function for YBX1. Such is the case during influenza virus infection, where binding of YBX1 to vRNPs facilitates their transport along microtubules from the perinuclear region into the vesicular trafficking system, which is required for the production of progeny virions (28). Similarly, during infection with the retroviruses human immunodeficiency virus (HIV) and mouse mammary tumor virus (MMTV), YBX1 is required for virus production. By promoting vRNA stability, YBX1 enables the formation of Gag-vRNA complexes, which are a pre-requisite for HIV assembly (29, 30). Furthermore, we previously reported that transient silencing of YBX1 expression in human Huh7 cells resulted in a decreased production of DENV infectious particles albeit with an increase in intracellular vRNA levels (9). Interestingly, YBX1 knockdown did not affect the percentage of infected cells, suggesting that the observed phenotype could be explained by an inhibition of vRNA packaging or virion egress (9).

Here we explore the function of YBX1 during DENV infection. We show that YBX1 localizes to DENV assembly sites and is required for the assembly of infectious virions after the formation of the nucleocapsid. Functional assays demonstrated that YBX1 enables the interaction between the viral proteins E and C, mediating the assembly of infectious virions and their secretion.

## RESULTS

### YBX1 is required for the production of DENV but is dispensable for the replication of intracellular vRNA

In order to understand the mechanistic basis for our previous studies (9) on the requirement of YBX1 for DENV productive infection, we abolished its expression in Huh7 cells by using CRISPR-Cas9 genomic editing. YBX1 ablation was confirmed by both genomic DNA sequencing (Fig. S1) and Western blot analysis (Fig. 1A). Three independently derived YBX1 KO clones (hereafter designed KO1, KO2 and KO3) were selected for further studies and all infectivity assays were conducted with DENV serotype 2. Consistent with our previous report (9), DENV-infected YBX1 KO cells showed significantly lower production of infectious viruses as well as significantly lower number of vRNA copies in supernatant compared to WT cells as determined at 18 and 24 hours post infection (hpi) (Fig. 1B – 1C). We also tested DENV dependency for YBX1 in the human lung epithelial A549 cell line and observed a significant reduction on DENV titers upon siRNA-mediated knock down of YBX1 (Fig. S2). These results confirm that in two different human-derived cell lines, both knockdown and knockout of YBX1 expression result in a significant decrease of progeny virus.

**Figure 1.**
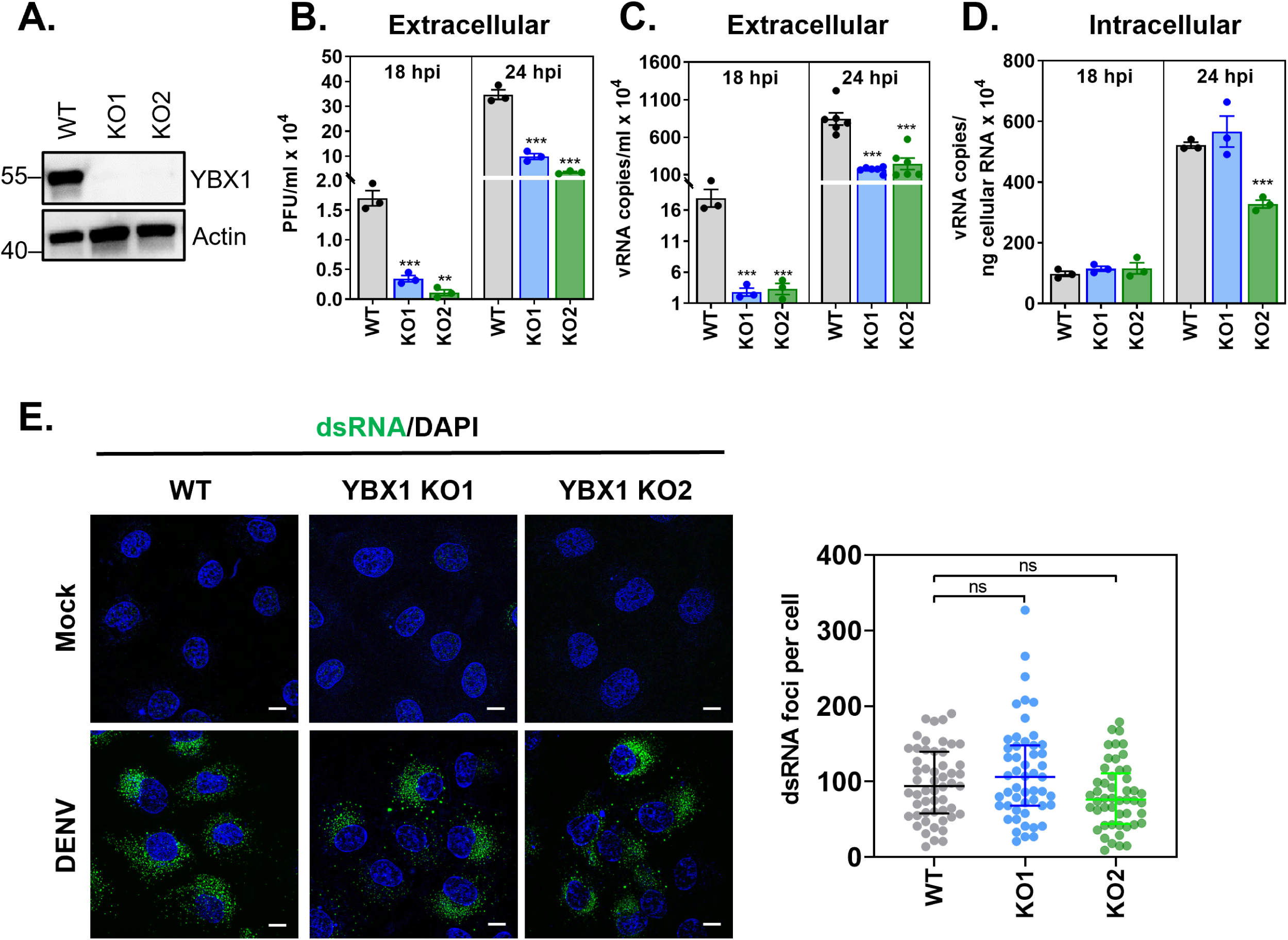
YBX1 is required for production of DENV progeny. **(A)** CRISPR/Cas9-derived YBX1 knockout (KO) clones (KO1 and KO2) do not express YBX1 protein as determined by western blot. **(A-E)** Huh7 wild type (WT) and YBX1 KO cells were infected with DENV at multiplicity of infection (MOI) of 1. **(B)** Supernatant virus titers were determined by plaque assay and expressed as plaque forming units (PFU) per ml. The number of vRNA copies was determined by RT-qPCR in supernatants **(C)** and intracellularly **(D)**. **(E)** Mock-infected and DENV-infected cells were processed for double stranded RNA (dsRNA) immunofluorescence at 24 hpi (green). Nuclei were stained with DAPI (blue). dsRNA foci per cell was determined in at least 30 cells per experiment using an in-house macro for ImageJ. Data are presented as mean ± SEM **(B-D)** and median ± IQR **(E)** from at least three independent experiments.

The reduction in DENV vRNA copies and infectious particles observed in supernatants from infected Huh7 YBX1 KO cells stands in sharp contrast with the intracellular vRNA levels, which were not affected at 18 hpi in either KO and were only affected in KO2 at 24 hpi (Fig. 1D). This result suggests that YBX1 is dispensable for the replication of DENV RNA genome. To further confirm that vRNA replication is not affected by YBX1 depletion, we evaluated the formation of double stranded RNA (dsRNA), which is a replication intermediate. No differences were observed between WT and YBX1 KO cells regarding the number of dsRNA foci per cell (Fig. 1E). These data confirm that genome replication is unaffected in YBX1 KO cells and suggest that this RBP is required at later stages of the replication cycle.

### YBX1 localizes to DENV assembly sites

As a first step to elucidate the function of YBX1 during DENV replication cycle, we studied its cellular localization upon infection. Whereas YBX1 distributes evenly in the cytoplasm of mock-infected cells, DENV infection causes the re-localization of YBX1 around the perinuclear region (Fig. 2A). DENV infection causes re-arrangements of the ER membranes, where replication of vRNA and assembly of new virions takes place at neighboring but distinct locations (4). In order to determine whether YBX1 localizes to DENV replication and/or assembly sites, we performed immunofluorescence co-staining of YBX1 with dsRNA, a marker of replication complexes, or with DENV E protein, which is enriched at the sites of virus assembly. YBX1 strongly colocalizes with E protein whereas limited colocalization was observed with dsRNA (Fig. 2B). The median percentage of signal intensities that overlapped with that of YBX1 was 70% (66-77) and 40% (30-45) for E and dsRNA, respectively. The overlap with YBX1 had a Mander’s coefficient of 0.71 (0.66-0.77) and 0.4 (0.3-0.45) and a middle Pearson’s coefficient of 0.5 (0.42-0.56) and 0.2 (0.09-.26) for E and dsRNA, respectively (Fig. 2C).

**Figure 2.**
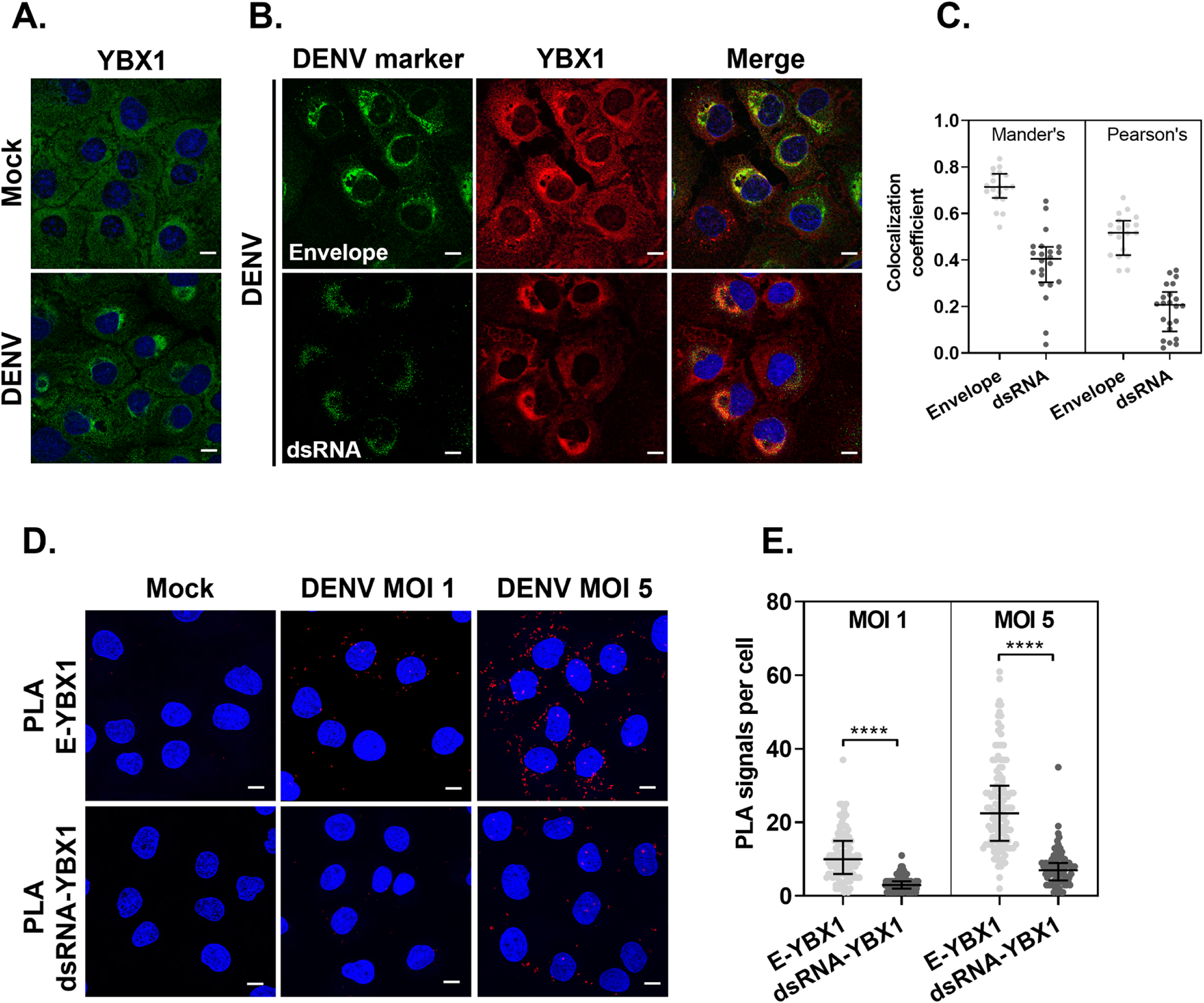
YBX1 localizes to DENV assembly sites. **(A-B)** Representative immunofluorescence of Huh7 cells mock-infected or infected with DENV at MOI 5 for 24 hpi and stained for the indicated markers. **(C)**Pearson’s and Manders’ correlation coefficients were extracted from the data set generated by the coloc2 plugin for imageJ. **(D)** At 24 hpi, Huh7 cells were processed for *in situ* proximity ligation assay (PLA) of E and YBX1 (upper panels) and dsRNA-YBX1 (lower panels). Nuclei were stained with DAPI (blue) and PLA signals are detected as red puncta. **(E)** The number of PLA signals per cell were counted in at least 30 cells per experiment using an in-house macro for ImageJ. Data are presented as median ± IQR from two independent experiments.

To further study the interaction between YBX1 and E proteins, we performed *in situ* proximity ligation assays (PLA), which has been used to detect protein-protein interactions with higher specificity and sensitivity compared to traditional co-immunostainings. PLA signals, detected as discrete fluorescent foci, depend on the recognition of target proteins (or dsRNA) by antibodies conjugated to oligonucleotide probes, which function as templates for rolling circle DNA amplification if the interacting factors targeted are within 40 nm of each other (31). In DENV-infected cells, E-YBX1 PLA signals were readily detected in a multiplicity of infection (MOI)-dependent manner (Fig. 2D). At MOI of 1, a median of 10 (6-15) E-YBX1 PLA foci were detected per cell, whereas 22.5 (15-30) E-YBX1 PLA foci were detected at MOI of 5 (Fig. 2E). Significantly lower number of dsRNA-YBX1 PLA foci were detected, with 3 (2-4) and 7 (4.2-9) foci per cell detected at MOI 1 and 5, respectively (Fig. 2E). These results suggest that upon DENV infection, YBX1 re-localizes to DENV assembly sites, where YBX1 is in close proximity to the structural viral protein E.

### YBX1 is required for the assembly of DENV particles after the formation of the nucleocapsid

Upon DENV vRNA replication, the structural viral protein capsid (C) binds vRNA to form the nucleocapsid, which is packaged into newly assembled virions. Given the re-localization of YBX1 to DENV particle assembly sites and its RNA-binding function, we hypothesized that interaction of this protein with DENV vRNA and/or C protein, would enable the loading and/or organization of vRNA during virus assembly. We tested the interaction of YBX1 with vRNA using RNA immunoprecipitation (IP). Lysates from DENV-infected WT Huh7 cells were immunoprecipitated using an IgG control or a YBX1 antibody. RNA was purified from the IP material and DENV vRNA copies were determined by RT-qPCR. Results presented in Fig. 3A show a significant 75 ± 16.6-fold enrichment of DENV vRNA copies in the YBX1-IP compared to the IgG-IP, confirming previous reports on the interaction of YBX1 with DENV vRNA (8, 9).

**Figure 3.**
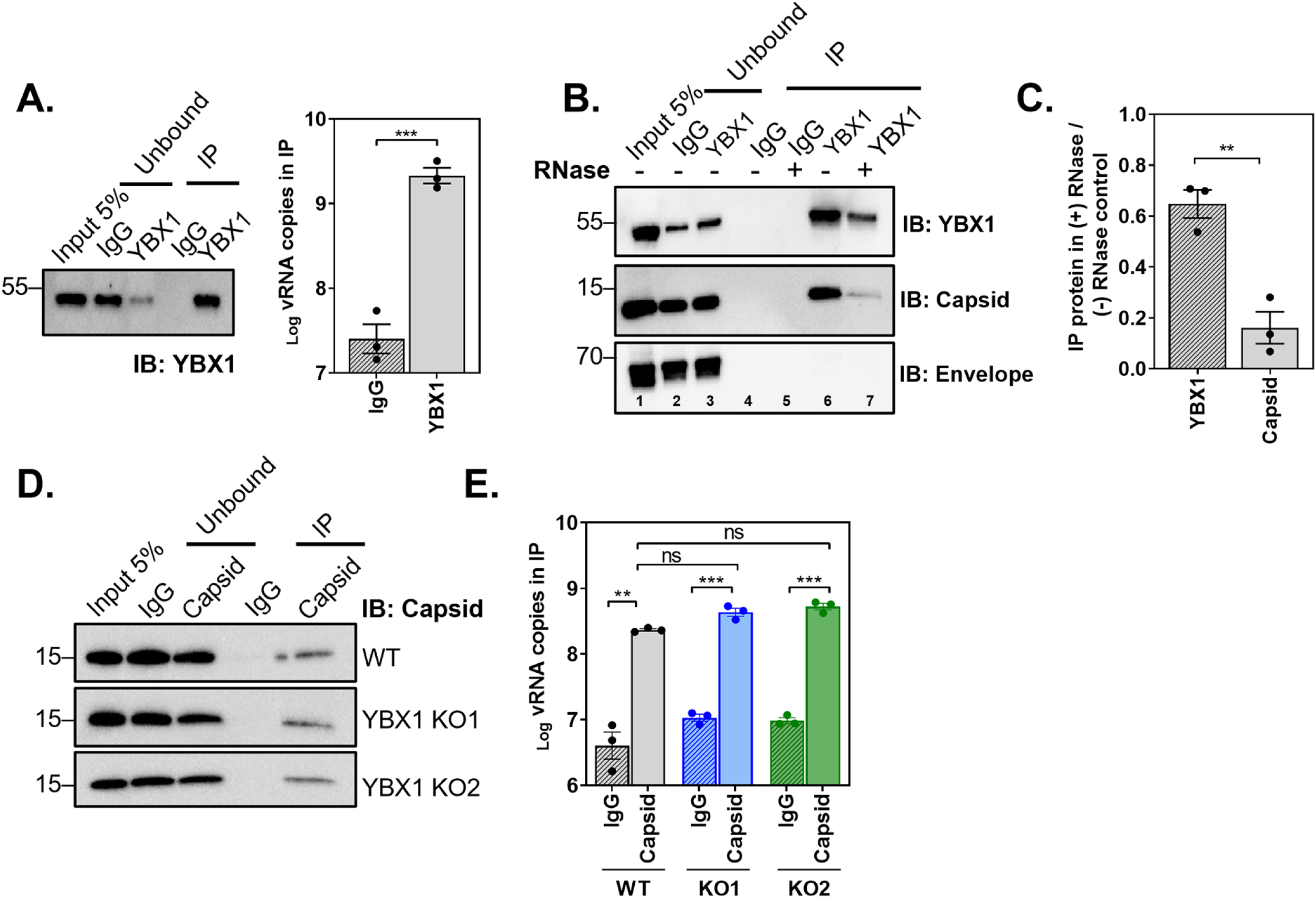
YBX1 interacts with DENV vRNA and capsid protein but is dispensable for nucleocapsid formation. Huh7 WT and YBX1 KO cells were infected with DENV at MOI 5 and processed for RNA-immunoprecipitation (RIP) or Co-IPs at 24 hpi. Cell lysates were incubated with an isotype control (IgG) and an antibody against YBX1 or capsid. Bound material was captured on Dynabeads protein G. **(A)** YBX1-RIP. Representative immunoblot of YBX1-pull down and total number of vRNA copies as determined in the IP by RT-qPCR. **(B)** IgG and YBX1-IPs were left untreated or treated with RNase A/T1 and immunoblots were performed for YBX1, capsid and envelope. **(C)** Densitometry analysis was performed to determine the levels of YBX1 and capsid in the IP sample treated with RNase relative to the non-treated control (lane 7 versus 6). **(D)** Capsid-RIP. Representative immunoblot of capsid-pull down and total number of vRNA copies as determined in the IP by RT-qPCR in WT and YBX1 KO cells. Data are presented as mean ± SEM from three independent experiments.

We next asked if YBX1 interacts with C and if such interaction is RNA mediated. To this end, we conducted co-IP experiments in the presence and absence of RNases A and T1 where IgG control or YBX1 antibody were used as baits for pull downs and immunoblots were stained for C. As shown in Fig. 3B, YBX1-IP effectively pulled down C protein (lane 6), suggesting that YBX1 and C interact in the context of infected cells. In contrast, E protein was not recovered in YBX1-IP samples despite the strong colocalization observed in Fig. 2. Importantly, RNase treatment decreased by 6.2 ± 0.94-fold the recovery of C protein in the YBX1 pull down (Fig. 3B, lane 7 and Fig. 3C), strongly suggesting that the YBX1-C interaction is RNA-mediated.

Given the interaction of YBX1 with DENV vRNA and C, we next investigated whether or not YBX1 was required for the interaction between vRNA and C and thus the formation of the nucleocapsid. We carried out RNA-IP experiments using lysates from DENV-infected WT and YBX1 KO cells and an IgG control or a C antibody (Fig. 3D). We did not observe differences in the number of vRNA copies recovered in the C-IP from WT and YBX1 KO cells (Fig. 3E). Our results indicate that YBX1 is not required for the formation of the nucleocapsid.

### YBX1 is required for the interaction between E and C proteins

Next, we investigated whether YBX1 could bridge the nucleocapsid to the assembly sites. We used PLA to study the interaction between E and C proteins in WT and YBX1 KO cells. E-C PLA signals were detected in DENV-infected WT cells with a median of 11 (7-19) foci per cell. A significantly lower association between E and C was observed in YBX1 KO1 and KO2 cells, with a median of 5 (2-8) and 7 (3-13) E-C PLA signals per cell, respectively (Fig. 4A). We also quantified the amount of E-prM PLA signals. We did not find a decrease of E-prM PLA signals per cell when comparing WT and YBX1 KO cells. WT cells had a median of 26 (18-54) per cell *versus* 58 (36-94) in KO1 and 27 (14-44) in KO2 cells (Fig. 4B). The difference observed between KO1 and KO2, could be explained by the intrinsic variability that arises from single-cell clonal expansion, but regardless we could conclude that YBX1 does not decrease the E-prM association. In contrast, a significant reduction in the interaction between E and C was consistent in the two YBX1 KO clones, suggesting that this RBP enables the association of the nucleocapsid with E.

**Figure 4.**
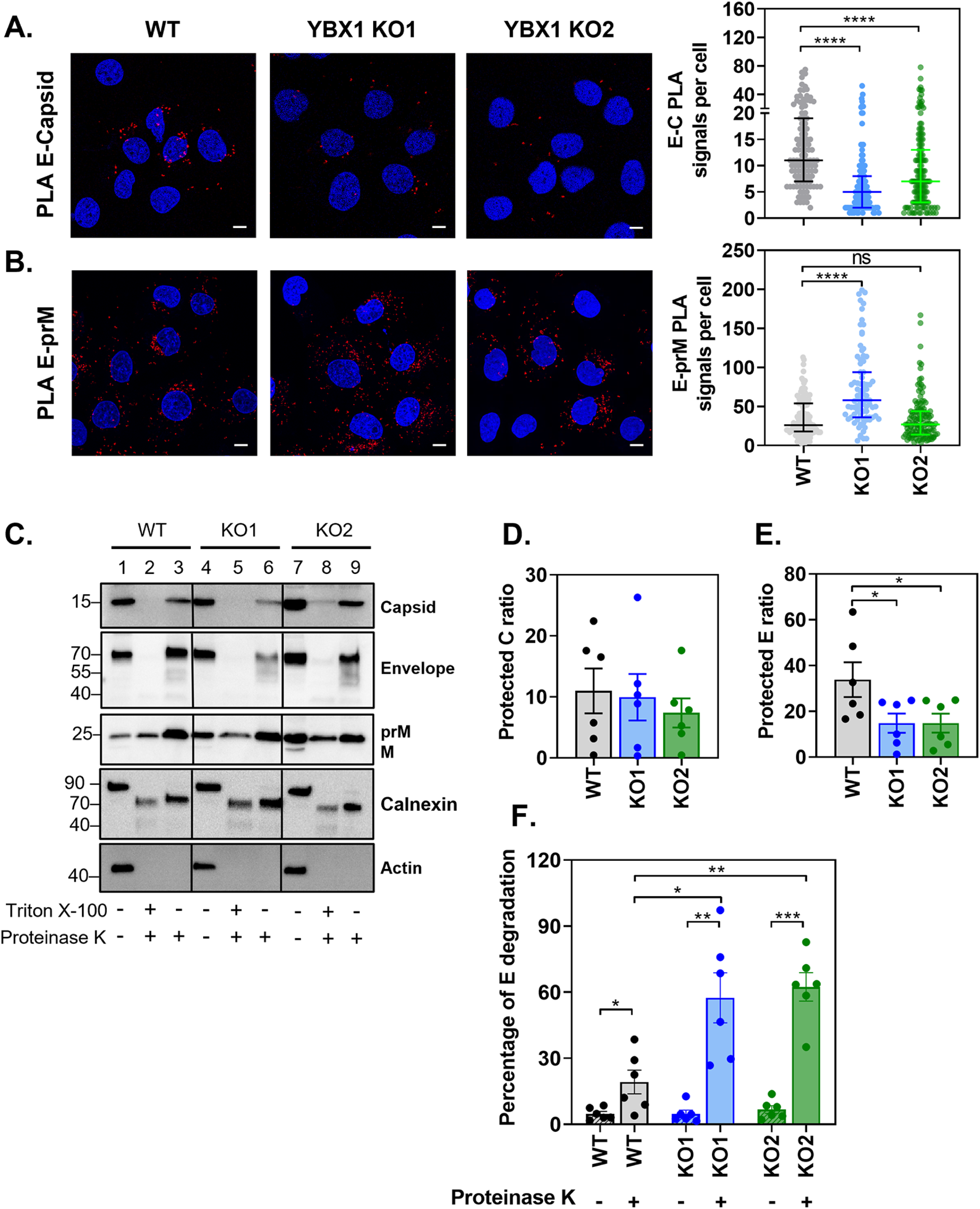
YBX1 promotes the interaction between E and C proteins. (A-B) **(A-B)** Huh7 WT and YBX1 KO cells were infected at MOI 5 and processed at 24 hpi for **(A)** E and capsid PLA and **(B)** and E and prM PLA. Nuclei were stained with DAPI (blue) and PLA signals are detected as red puncta. The number of PLA signals per cell were counted in at least 50 cells per experiment using an in-house macro for ImageJ. Data are presented as median ± IQR from two independent experiments. **(C-F)** Proteinase K (PK) protection assay of protein lysates from DENV-infected WT and YBX1-KO cells. Equal amounts of protein were left untreated or treated with PK in the absence and presence of triton X-100 and samples were analyzed by western blot **(C)**. The ratio of protected E protein **(D)** and C protein **(E)** was calculated by dividing the densitometry values of the PK-treated samples by that of the non-treated control. The percentage of E protein degradation **(F)** was calculated by dividing the densitometry value of the 70 KDa band by the densitometry value of the entire lane. Data are presented as mean ± SEM from six independent experiments.

To further probe this idea, we adapted a previously described proteinase K (PK) protection assay to measure the envelopment of C protein (32). In fully assembled enveloped viruses, the nucleocapsid is surrounded by a lipid bilayer and hence resistant to the action of proteases; however, pre-treatment with detergents, which remove ER and viral membranes, expose the capsid viral protein to the action of proteases. We reasoned that if the nucleocapsid packaging (envelopment) depended on YBX1, we would observe greater degradation of C protein by PK in the absence of detergent in YBX1 KO cells relative to WT cells. Consequently, lysates from WT and YBX1 KO cells were treated with PK in the presence and absence of triton X-100 and the levels of structural viral proteins were determined by western blot (Fig. 4C). We quantified the ratio of protein protected from PK-degradation by dividing the densitometry values of the PK-treated samples by that of the non-treated control. We found a slight reduction of the fraction of C that was protected from PK degradation between WT and YBX1 KO cells; however, this difference was not statistically significant (Fig. 4C, lanes 3, 6 and 9, and Fig. 4D). Although this assay suggested that C is equally protected from PK-mediated degradation in all cell lines, the noisy quality of the data did not unambiguously rule out an effect of YBX1 in envelopment.

These experiments, however, resulted in an unexpected observation: in YBX1 KO cells, E protein was more labile to PK-mediated degradation as shown by the reduced levels of protected E ratio (Fig. 4E). In PK-treated samples, there was a reduction in the intensity of the E band, which is detected at 70 KDa, and the appearance of partial degradation products of E protein (Figure 4C, lanes 3, 6, 9).

We quantified the levels of E protein degradation by dividing the densitometry value of the 70 KDa band by the densitometry value of the entire lane. As shown in Fig. 4F, in WT cells, 19.19% ± 5.39 of E protein is degraded following treatment with PK. In contrast, in YBX1-KO cells a significantly higher percentage of E degradation is observed (57.43 ± 11.37% for KO1 and 62.40 ± 6.45 % for KO2). This effect was specific for E, because no differences were observed on the degradation of prM, nor the host ER-resident protein calnexin (Fig. 4C). Our results indicate that ablation of YBX1 reduces the interaction between E and C proteins and increases the susceptibility of E to PK-mediated degradation.

### YBX1 is required for the assembly of intracellular infectious virions

The reduced interaction between E and C proteins and the higher susceptibility of E to PK-mediated degradation observed in YBX1 KO cells suggested that virus particles were not properly assembled. To test this, we analyzed WT and YBX1 KO DENV-infected cells via transmission electron microscopy (TEM). DENV infection causes drastic rearrangements of the ER membrane, which are readily identified under TEM as convoluted membranes and vesicle packets (Vp). In addition, intracellular DENV virions are detected in the lumen of the ER as electron dense particles with a diameter of around 45 nm (4). Indeed, in WT DENV infected cells, we observed convoluted membranes, Vps and viral particles (Vi) (Fig. 5A). Vps, the sites of viral RNA replication, are observed in close proximity to assembled virions, which were detected in arrays as well as individual particles in the lumen of the ER (Fig. 5A). In YBX1 KO cells, Vps were also detected and no differences regarding their size or amount were apparent compared to WT cells (Fig. 5B – C). This is consistent, with normal levels of dsRNA replication, which we found to be unaffected in YBX1 KO cells (Fig. 1E). Remarkably, however, in YBX1 KO cells we very rarely observed the typical electron dense DENV virions. Instead, we observed empty-looking virions, with a rough appearance and variable size in the ER lumen of the two YBX1 KOs examined, KO1 and KO3 (Vi_(KO)_) (Fig. 5B – C). Rarely we noted that these empty-looking virions appeared to be attached to the ER membrane (Fig. 5B, left panel).

**Figure 5.**
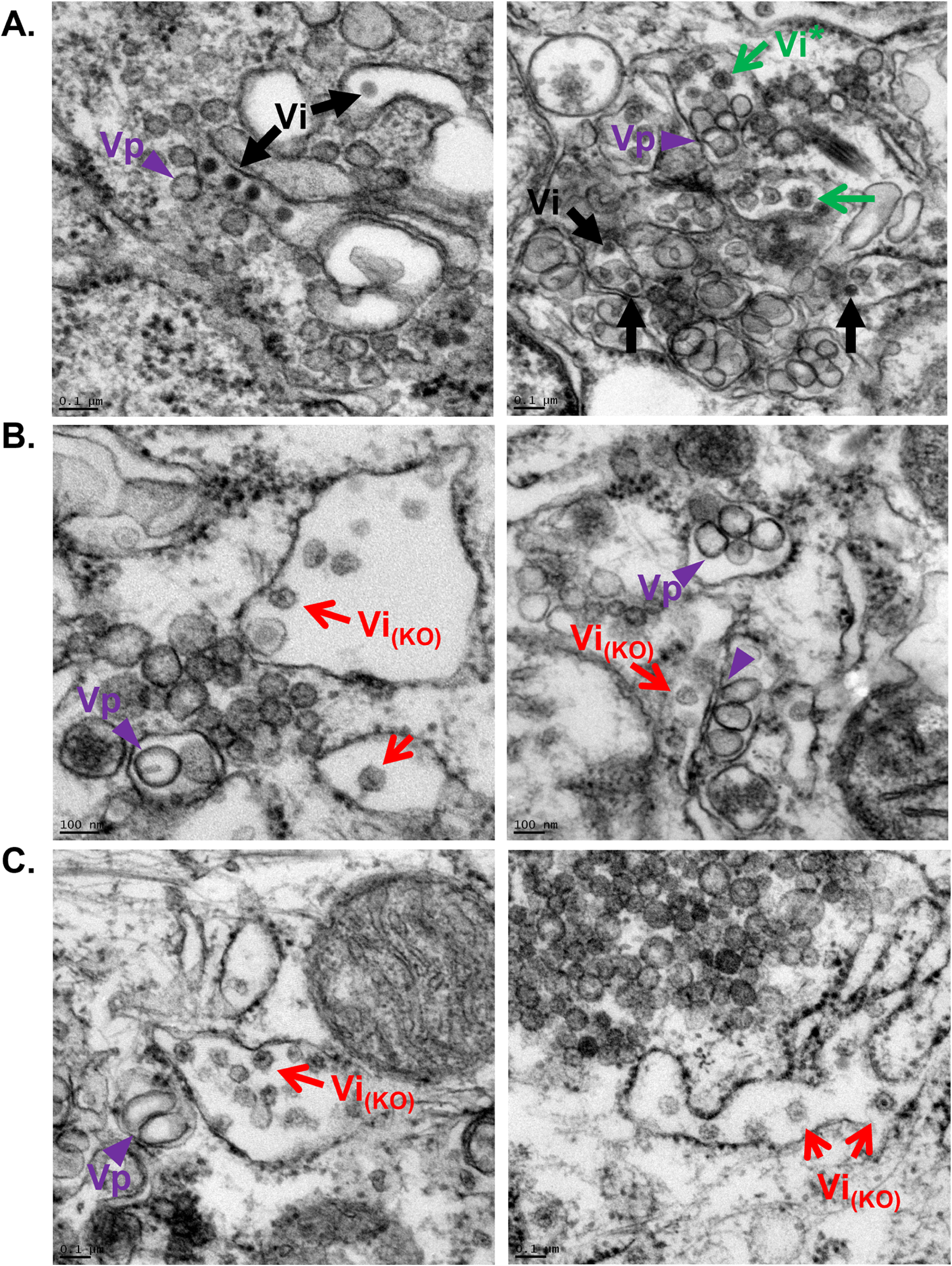
Thin section TEM images of DENV-infected resin-embedded WT **(A)** and YBXI KO **(B-C)** Huh7 cells. Virus-induced structures include vesicle packets (Vp, purple arrow heads), which are the site of viral RNA replication. DENV infectious virions are identified as electron dense particles (Vi, black arrows) in arrays (A, left panel) and as individual virions (A, right panel). Empty looking particles with rough surface are detected in high abundance in the ER lumen of YBX1 KO cells (red arrows) and sporadically in WT cells (green arrows).

Furthermore, an electron dense region was observed in the center of some of these rough particles (Fig. 5B, left panel, Fig. 5C). Interestingly, rough virions with the same electron dense dot were seen sporadically in WT cells (Vi* in Fig. 5A, right panel), which is consistent with the fact that not all virions produced by infected cells are properly assembled (33). Lastly, the empty-looking rough virions in the ER of KO cells appeared to be more frequent than virions in WT cells, suggesting that these particles are not efficiently secreted to the extracellular milieu.

The empty-looking aspect of the particles observed in YBX1 KO cells, suggested that they are not infectious particles. Therefore, we next determined whether the infectivity of cell-associated virus particles was affected in YBX1 KO cells. To complete the processing of assembly, DENV virions bud into the ER lumen, thereby acquiring a lipid membrane that contains heterodimers of the structural proteins E and prM. These newly assembled immature virions traffic through the secretory pathway and, in the trans Golgi network, the cellular protease furin cleaves the pr-peptide from the M protein (34). This processing is essential to the formation of mature infectious particles. In order to test the requirement of YBX1 for the formation of infectious particles independently of their maturation status, recovered intracellular virus was left untreated or incubated with furin (Fig. 6A). The results in Fig. 6B show that significantly lower intracellular virus titers are detected in YBX1 KO cells.

**Figure 6.**
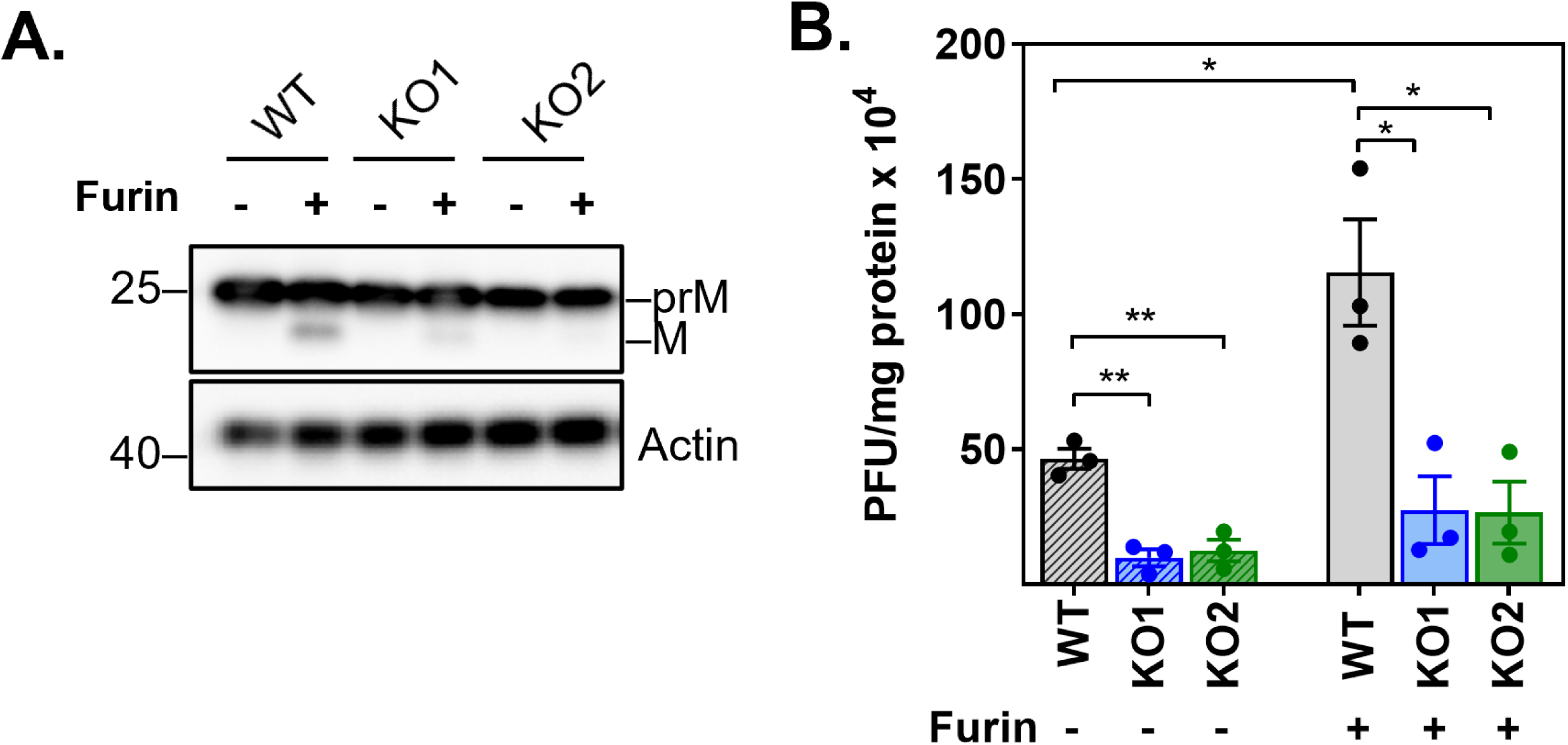
YBX1 is required for the assembly of intracellular infectious virions. Huh7 WT and YBX1 KO cells were infected with DENV at MOI 5. At 24 hpi, cell pellets were harvested and subjected to 5 cycles of freeze and thaw. Equal amounts of protein lysates were left untreated or treated with furin for 16 h at 30 ℃. **(A)** Representative western blot analysis of prM and M expression upon furin treatment. **(B)** Titration of protein lysates was determined by plaque assay. Titers are expressed as PFU per milligrams of protein. Data are presented as mean ± SEM from three independent experiments.

Compared to WT cells, YBX1 KO1 and KO2 cells show a 6.56 ± 2.9 and 4.7 ± 1.69-fold reduction in cell-associated infectious virus, respectively. As expected, treatment with furin increased the intracellular virus recovered from WT-infected cells. Furin treatment, however, did not rescue the YBX1 requirement as accumulation of intracellular virus in YBX1 KO1 and KO2 cells was 5.97 ± 1.99 and 6.24 ± 2.35-fold less than in WT cells. These data indicate that YBX1 does not influence virion maturation, but it is required for the assembly of infectious viruses.

### YBX1 is required for efficient secretion of virions

In addition to infectious virus, DENV-infected cells secrete virus-like particles (VLPs), which contain E and prM but are non-infectious because they lack the nucleocapsid (35–37). Therefore, defects in the envelopment of the nucleocapsid do not necessarily translate in lower particle secretion. Because we observed an apparent accumulation of abnormal virions in the ER lumen of YBX1 KO cells (Fig. 5B – C), we investigated whether particle secretion was also affected in these cells. To achieve this goal, we compared the intracellular and extracellular accumulation of E and C proteins between WT and YBX1 KO cells. Whereas no significant differences were observed in the intracellular levels of E and C proteins between WT and YBX1 KO cells, a strikingly lower accumulation of both proteins was observed in the media from YBX1 KO cell cultures (Fig. 7A – C). E and C expression were reduced by 4.48 ± 0.95 and 12.05 ± 0.95 fold in YBX1 KO1 cells and 6.68 ± 0.98 and 35.29 ± 0.97 fold in YBX1 KO2 cells, respectively. In addition, we also found that compared to WT cells, YBX1 KO cells secrete lower amounts of the non-structural viral protein NS1 (Fig. 7D). NS1 localizes to the ER lumen and, interestingly, it has been shown to interact with E and prM to assist in the budding step (38). Altogether these data show that secretion of E protein (a proxy of VLPs) and NS1 is affected by YBX1 depletion, strongly indicating that YBX1 is required for the secretion of DENV particles.

**Figure 7.**
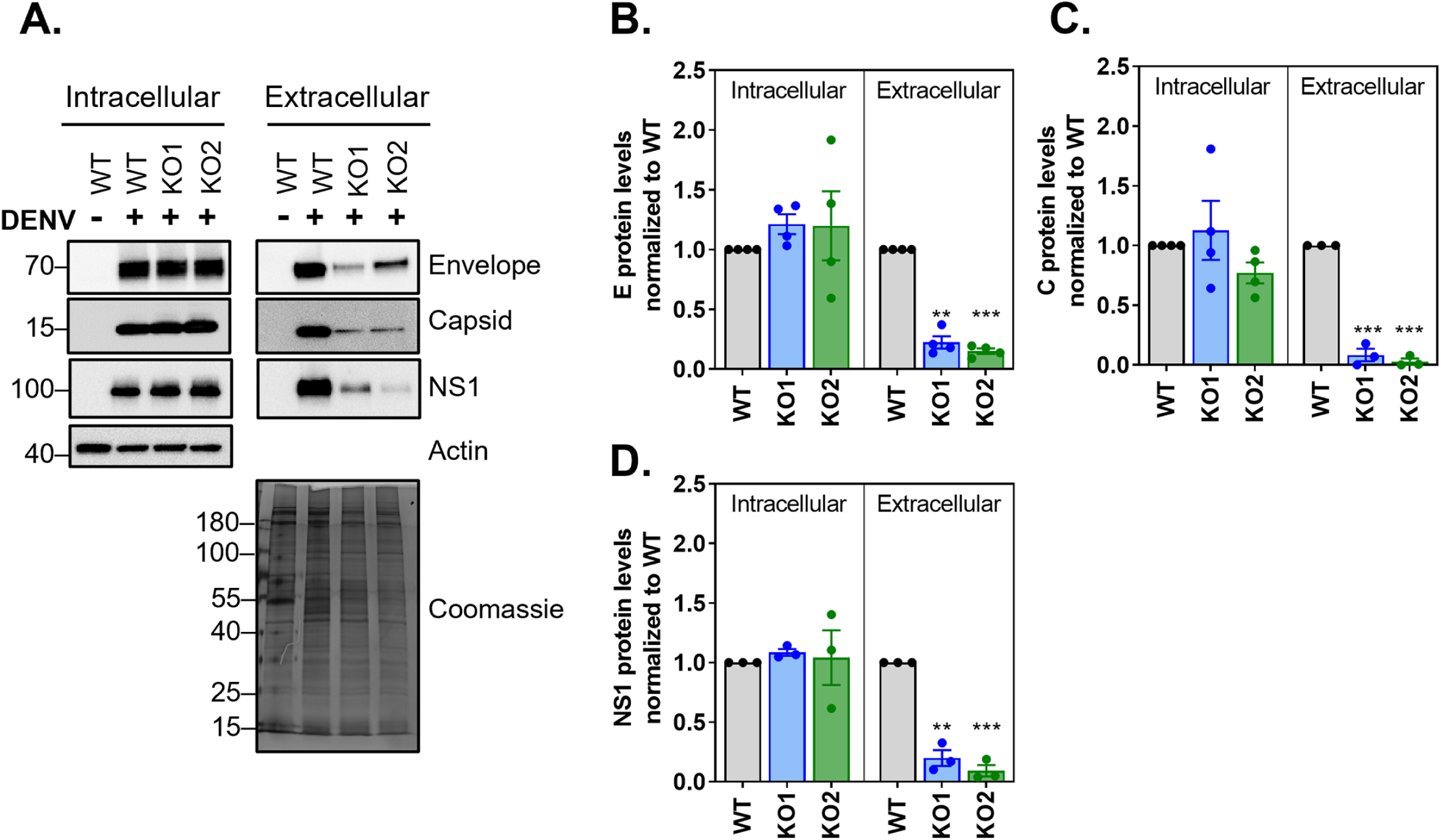
YBX1 is required for viral secretion. **(A)** Huh7 WT and YBX1 KO cells were infected at MOI 5 and cell pellets and supernatants were analyzed via western blot at 18 hpi. **(B-D)** The intracellular levels of the indicated viral proteins was normalized to that of actin and this value was used to normalize the levels detected in supernatant for each protein. In all cases, the expression relative to WT cells is shown. Data show mean ± SEM from at least three independent experiments.

## DISCUSSION

The assembly of *Flavivirus* particles involves a series of events tightly coordinated in time and space. Upon vRNA synthesis, single vRNA molecules exit from the replication complexes towards vicinal assembly sites. Coupling of vRNA with C proteins gives rise to the nucleocapsid, which is enveloped by E and prM protein-containing membranes as newly assembled particles bud into the lumen of the ER (Fig. 8). These virions are transported via the secretory pathway and reach the extracellular milieu by exocytosis. Here, we report that the host RBP YBX1 is required for the production of DENV by assisting at the assembly step and promoting the egress of virus particles. We show that YBX1 interacts with vRNA and capsid but is dispensable for vRNA replication and formation of the nucleocapsid. In contrast, ablation of YBX1 decreased the spatial proximity between E and C proteins, which resulted in the assembly of viral-like particles with low electron density in TEM and lower DENV yields. Although we cannot formally rule out indirect effects mediated by alterations of host gene expression by YBX1 depletetion, physical interactions of YBX1 with vRNA and capsid, and YBX1 localization to DENV assembly sites favor a direct role for YBX1 on DENV infection.

**Figure 8.**
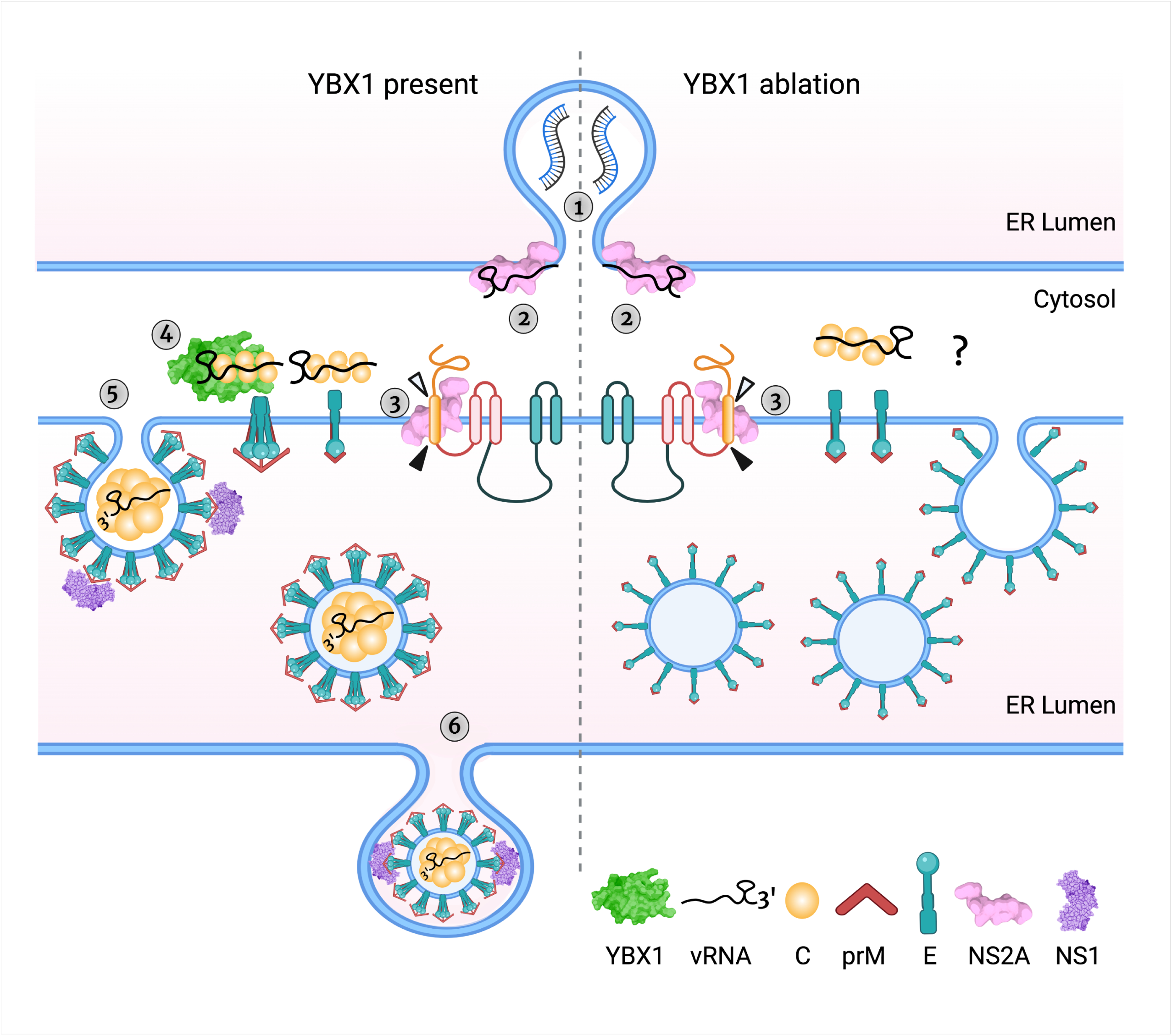
YBX1 is required for DENV budding: a model. The schematic shows DENV assembly and egress steps in WT (left of the dashed line) and YBX1 KO cells (right of the dashed line) (1) There is no difference in vRNA accumulation in WT and YBX1 KO cells and we interpret this to indicate normal DENV vRNA replication, which occurs in ER invaginations in close proximity to the assembly sites. (2) NS2A binds to vRNA and recruits it to the assembly sites in both WT and KO cells. (3) Processing of capsid-prM junctions allows the formation of prM-E dimers, the dimerization of capsid proteins and its association with vRNA. (4) YBX1 binds to the nucleocapsid enabling the interaction of capsid dimers with prM-E trimers. We posit that this interaction is disrupted in YBX1 KO cells. (5) The viral membrane proteins drive the budding of assembled particles into the ER lumen. In YBX1 KO cells, budded particles are empty-looking and with a rough surface. We propose that these anomalous particles arise from an assembly defect affecting the organization of E on the host membrane, likely due to the lack of interaction between this protein with the nucleocapsid, which is mediated by YBX1. (6) Assembled particles are secreted via the secretory pathway, in a process likely mediated by NS1 and other sorting mechanism involving prM-E interacting proteins. In the anomalous particles found in YBX1 KO cells, the secretion motifs are lacking or hidden, and thus particles accumulate in the ER lumen. Created with BioRender.com

The demonstration that DENV yield is decreased in YBX1 KO Huh7 cells and A549 cells where YBX1 was knocked down is in line with our previous report demonstrating that transient silencing of YBX1 via siRNAs resulted in lower extracellular DENV virus titers (9). Consistent with this, siRNA-mediated knockdown of YBX1 in Huh7 cells resulted in a significantly lower production of Zika virus, a *Flavivirus* closely related to DENV, despite limited effect on intracellular viral protein expression and vRNA (39). In contrast, using YBX1 KO MEFs, Paranjape and Harris showed that ablation of YBX1 promoted infectious virus production (8). This paradoxical result can be explained by the fact that YBX1 KO mice exhibit embryonic lethality (E13.5) (40, 41) and fibroblasts derived from YBX1 KO embryos show diminished oxidative stress responses (41). In addition, YBX1 KO zebrafish embryos display elevated levels of global translation and an activated unfolded protein response (42). Given that oxidative stress responses have been shown to mediate antiviral innate immune responses to DENV (43) and that activation of the unfolded protein response is required for productive infection (44, 45), it is likely that the anomalous phenotype exhibited by YBX1 KO MEFs is *per se* conducive to DENV infection and thus, the proviral function of YBX1 was masked in these cells. From our work on DENV and that of Bonenfant and colleagues (39), we conclude that efficient *Flavivirus* assembly requires YBX1 in human cells.

Interestingly, two other host RNA-binding proteins, Nucleolin and DDX56, are required for the assembly of DENV and WNV, respectively (46–48). Although their precise function during assembly is not well understood, both proteins were shown to interact with capsid in an RNA-independent manner. Conversely, we found that YBX1 interactions with C protein were RNA-dependent, likely mediated by vRNA, given that we and others have shown direct interaction of YBX1 with DENV vRNA (8, 9). In fact, previous studies revealed that YBX1 binds to DENV 3’ stem loop, a highly conserved region among Flaviviruses located at the 3’ end of the 3’UTR (8).

Recent studies highlight the importance of the 3’UTR for the completion of DENV and ZIKV assembly (49, 50). Xie and collaborators demonstrated that the non-structural protein NS2A binds to DENV 3’UTR to recruit newly synthesized vRNA from the replication complex towards the assembly sites. In addition, separate NS2A molecules interact with the nascent C-prM-E polyprotein and the viral protease.

Oligomerization of NS2A molecules brings together the newly synthesized vRNA and the C-prM-E polyprotein, leading to the coordinated cleavage of C from prM-E, the formation of the nucleocapsid and its envelopment (49). We propose here to include YBX1 as an essential component of these coordinated molecular events (Fig. 8), given that our results indicate that by interacting with the nucleocapsid, YBX1 promotes the proper assembly of infectious particles. Several lines of evidence support this idea; first, YBX1 colocalized with E and interacted with C protein in an RNA-dependent manner. Second, ablation of YBX1 weakened the interaction between E and C protein and resulted in the formation of empty-looking particles, consequently reducing the intracellular virus titers. It remains to be investigated whether YBX1 interacts with NS2A and/or the NS2A-vRNA complex and the precise timeline of YBX1-vRNA interactions. Nevertheless, since we did not observe defects in processing of the viral structural proteins in DENV-infected YBX1 KO cells and the formation of the nucleocapsid was not affected, we hypothesize that YBX1 is dispensable for the transport of vRNA by NS2A to the assembly sites and for the processing of the C-prM-E polyprotein. Instead, YBX1 is likely to bind the nucleocapsid once it has been formed to provide molecular clues important for subsequent steps in virion assembly (Fig. 8).

On the one hand, the PK protection assay suggested that envelopment of the nucleocapsid was not affected by YBX1 KO, but the data were not conclusive (Fig. 4C). On the other hand the TEM studies showed that most of the viral like particles formed in YBX1 KO cells had low electron density and rough surfaces (Fig. 5) resembling the DENV2 and TBEV VLPs formed in the absence of nucleocapsid (51–53). Further supporting a defect in envelopment are the data showing disrupted E-C interactions in KO cells (Fig. 4A). These abnormally assembled viruses are not infectious (Fig. 6). We propose that in YBX1 KO cells DENV nucleocapsids are enveloped inefficiently (Fig. 8).

As noted above the empty looking viral particles observed in YBX1 KO cells have a rough appearance suggesting that the E protein on their surface is not folded as seen in virions in WT cells. This is consistent with the increased sensitivity of E to PK treatment (Fig. 4C, E & F). All of these findings suggest that in the absence of YBX1 E proteins are not organized properly on the virion (Fig. 8).

We further propose that abnormal assembly of DENV leads to defects in virion secretion and explains the lower levels of extracellular E protein observed in DENV-infected YBX1 KO cells (Fig. 7 & 8). This contrasts with VLPs, which are readily secreted after the expression of only E and prM proteins (35–37). The empty-looking rough particles seen in YBX1 KO cells vary in size but many are larger than virions, whereas VLPs are usually significantly smaller than infectious virions (Fig. 5), which range from 25 to 31nM (51–53). Interestingly, we also observed lower secretion of NS1 from DENV-infected YBX1 KO cells. The interaction of NS1 with E and prM, which has been shown to mediate membrane bending and envelopment of nucleocapsids, is critical for the production of infectious DENV (38). In addition, NS1 is required for the initial vesicular trafficking of assembled virus particles (38).

Therefore, we posit that in the viral structures formed in YBX1 KO cells, E proteins poorly interact with NS1. Alternatively, in these anomalous viral particles, E and prM domains important for secretion could be inaccessible. For instance, the trimming of the N-linked glycan at position Asn154 of TBEV E protein has been shown to be crucial for the secretion of VLPs (52). In addition, it has been shown that interaction of KDEL receptors, which traffic between the ER and Golgi apparatus, and prM mediate vesicular transport of DENV VLPs, arguing that particle release is assisted by host factors that interact with viral surface proteins (54).

The mechanisms leading to the envelopment of *Flavivirus* nucleocapsids are largely unknown. Important questions that remain open is how viral RNA is recognized by the capsid protein, given that no packaging signals have been identified in the RNA, and how are the nucleocapsids enveloped assuring that assembled virions *only* contain viral RNA and proteins. Furthermore, knowledge regarding the function of cellular RNA-binding proteins in these processes is lacking. Here we report that the host protein YBX1 is required for the assembly of infectious virions, likely by enabling the sorting of the nucleocapsids at the budding sites.

Interestingly, the inward membrane budding (away from the cytoplasmic face) that leads to the assembly of new DENV virions evokes a range of cellular processes, such as the loading of cargo into exosomes and cytokinesis, which are mediated by the endosomal sorting complex required for transport (ESCRT) (55). Intriguingly, a role for YBX1 during these processes has been reported (22–25, 56). Particularly, accumulating evidence supports the requirement of YBX1 for the sorting of RNA cargo, mainly small non-coding RNAs, into exosomes (22–25). Because YBX1 does not contain transmembrane domains, it is likely that the intersection with exosomes that concludes with the loading of specific RNA cargo is mediated by YBX1-interacting partners. Indeed, previous studies have shown interaction between ESCRT proteins and YBX1 in the context of RNA or YBX1 secretion (57, 58).

Although the exact details of the mechanism by which YBX1 mediates productive DENV assembly remains to be unraveled, the results presented here provide important insights on the function of this RNA binding protein and we present a model that deepens our understanding of the late stages of DENV replication cycle.

## Acknowledgements

We thank members of the Garcia-Blanco-Bradrick Laboratory (UTMB) and Garcia-Blanco-Pompon Laboratory (Duke-NUS) for their insightful suggestions and support. We also thank Ms. Zhixia Ding for her expert assistance in TEM. We also would like to acknowledge Dr. Michael Sheetz and Dr. Pei-Yong Shi (UTMB) for their suggestions to the manuscript. MDT was the recipient of a Rubicon fellowship from The Dutch Research Council (NWO). The project was supported by the Duke-NUS Signature Research Programme in Emerging Infectious Diseases funded by the Agency for Science Technology and Research (A*Star), and NIH/NIAID P01 AI150585 to M.A.G-B, and NMRC (Singapore) grant NMRC/ZRRF/0007/2017 to J.P. The funders had no participation in the study design, data collection and analysis, decision to publish, or preparation of the manuscript.

## Supplementary Figure Legends

**Table S1. List of antibodies used.** WB: western blot; RNA-IP: RNA immunoprecipitation; IF: immunofluorescence; PLA:proximity ligation assay.

**Figure S1. CRISPR/Cas9-mediated genomic editing of YBX1 in KO1, KO2 and KO3.** Total DNA was extracted from Wild type (WT) Huh7 cells and three independent YBX1 knockout clonal cell lines (KO1, KO2 and KO3). The CRISPR/Cas9-targeted region (exon 3 of YBX1) was amplified by PCR, cloned into the pCRTM2.1-TOPO vector and used to transform DH5-α competent E. coli cells. Two recombinant clones (C1 and C2) per cell line were sequenced. The 20-base pair target sequence (sgRNA) is shown in green and the PAM sequence is shown in red.

**Figure S2. siRNA-mediated knockdown of YBX1 in A549 cells reduces DENV yield.** A549 cells were left non-transfected (NT), transfected with a non-targeting negative control siRNA (siNC) or two siRNAs targeting YBX1 (siYBX1#1 and siYBX#6) for 48 h prior to infection with DENV at MOI 1. **(A)** YBX1 and actin expression was assessed by Western blot at 48 h post transfection. (B) Viral titers were determined via plaque assay at 24 hpi. Data are presented as mean ± SEM from three independent experiments.

## Materials and Methods

### Cell lines, reagents and services

The human hepatic Huh7 cells (JCRB0403) were maintained in Dulbecco’s modified eagle medium (DMEM, Gibco, the Netherlands) containing 10% fetal bovine serum (FBS), 100 U/ml penicillin and 100 µg/ml streptomycin. BHK-21 cells (ATCC CCL-10) were grown in RPMI (Gibco) media supplemented with 10% FBS, 100 U/ml penicillin and 100 µg/ml streptomycin. All cells were cultured at 37℃ and 5% CO_2_. All sequencing services were carried out by First Base and all DNA oligos were ordered from Integrated DNA Technologies (IDT, Singapore). For transmission electron microscopy experiments Human hepatoma (HuH-7) wild type and YBX-1 KO cells were maintained in Dulbecco’s modified Eagle’s medium (DMEM, Genesee Scientific, San Diego, CA) supplemented with 10% fetal bovine serum (FBS, Genesee Scientific, San Diego, CA) and 100 U/ml penicillin and 100 μg/ml streptomycin (P/S, Gibco, Waltham, MA) at 37°C with 5% CO_2._

### Virus stock, titration assays and furin treatment

DENV serotype 2 strain New Guinea C was a kind gift from Aravinda de Silva (University of North Carolina, Chapel Hill, NC) and propagated in C6/36 cells as described before(9). The same virus stock was used for all experiments. Virus stocks and experimental samples were titrated by plaque assay. Briefly, BHK-21 cells were seeded in 24-well plates at a cell density of 6.5x10^4^ cells/well. At 24 h post-seeding, cells were infected with a 5-fold serial dilution of sample. Cells were infected for 1 h and an overlay of 1% carboxymethylcellulose (Aquacide II, Calbiochem, San Diego, CA) prepared in RPMI was added. Plaques were fixed and stained at 5 days post-infection and titers are reported as plaque forming units per ml (PFU/ml). To determine intracellular virus titers, virus-containing supernatant was removed and cells were washed thoroughly with plain media. Cells were scraped into DMEM containing 5% FBS and 15 mM HEPES (Gibco) and centrifuged at 300 g for 10 minutes at 4℃. Cell pellets were resuspended in 100 µl media and disrupted by 5 cycles of freeze/thaw using a dry ice/ethanol bath for freezing and a water bath set at 37℃ for thawing. Cell debris was removed by centrifugation at 2000 g for 15 min at 4℃. When indicated, cell lysates were adjusted to the same protein concentration using furin cleavage buffer (100 mM Tris-HCL and 1mM CaCl_2_) and incubated with 5 units of Furin (NEB, Ipswich, MA) for 16 h at 30℃. Virus titers were determined by plaque assay as described above.

### DENV infection

Cells were incubated for 1h at 37℃ with the indicated multiplicity of infection (MOI) of DENV-2 NGC and then washed three times. Cells were cultured further in DMEM media supplemented with 5% FBS and 15 Mm HEPES. Supernatants and cell lysates were harvested at the indicated time points.

### Absolute quantification of supernatant and intracellular vRNA copies

Extraction of vRNA from supernatants was performed using a QIAamp viral RNA mini kit (Qiagen, Hilden, Germany) following manufacturer’s instructions. Total RNA from cell pellets was extracted using E.Z.N.A. Total RNA kit I (OMEGA Bio-Tek, Norcross, GA), following manufacturer’s instructions. Concentration of total intracellular RNA was determined via Nanodrop (Thermo Fisher Scientific, Waltham, MA). For absolute quantification of vRNA, a one-step RT-qPCR reaction with iTaq Universal probe kit (Bio-Rad, Hercules, CA) and primers and probes targeting the DENV2 envelope were used (59). Absolute copy numbers were determined using a standard curve generated with an *in vitro* transcribed RNA fragment containing the qPCR target as detailed by Pompon et al (59). PCR reactions were conducted on a CFX96 Touch Real-Time PCR Detection System (Bio-Rad).

### Plasmids

The cloning plasmid pSpCas9-BB-2A-GFP was obtained from Addgene and the sgRNAs targeting the YBX1 gene were designed by the online CRISPR tool from the Broad Institute(60). We cloned the sgRNA 5’-GTCTTGCAGGAATGACACCA-3’, which targets exon 3 from YBX1 gene, into the pSpCas9-BB-2A-GFP vector following the protocol outlined in step # 5 (preparation of expression construct) from Ran *et al* (61). The generated construct pSpCas9-BB-2A-GFP-sgYBX1 was propagated in DH5-α competent *E. coli* cells (NEB) and validated by sequencing using the universal primer U6-Forward.

### CRISPR/Cas9-mediated knockout

Huh7 cells were seeded at a cell density of 2.3x10^5^ in 6-well plates 24 hours prior transfection. Transfection of 2 µg pSpCas9-BB-2A-GFP-sgYBX1 was carried out with 6 µl of Lipofectamine 2000 (Thermo Fisher Scientific) following manufacturer’s instructions. At 24 h post transfection, cells were sorted on a BD FACSAria II flow cytometer on the basis of GFP expression. Isolation of single cells from the GFP-positive population was carried out by serial dilution as suggested by Ran and collaborators(61). Plates were inspected carefully and those in which multiple cells were seeded were disregarded for following culture. Cells were cultured for 3-4 weeks to allow clonal expansion of knock out (KO) cells. KO of YBX1 gene was validated by western blot analysis and sequencing. Briefly, genomic DNA from wild type (WT) and YBX1 KO cells was extracted using DNeasy Blood & Tissue Kit (Qiagen) and amplified using the primers 5’-GGGTAATGAGGCTACAACTGTTT-3’ and 5’-GTCAACTCTAACACATTTCCACGTAT-3’. The PCR product was cloned into pCR^TM^2.1-TOPO vector (Invitrogen) and used to transform DH5-α competent *E. coli* cells (NEB). Recombinant clones were sequenced using the universal primers M13-Forward and M13-Reverse.

### Transfection of siRNAs

Two independent siRNAs targeting YBX1 (Hs_YBX1-1 and Hs_YBX1-6 FlexiTube siRNA, Qiagen) and a non-targeting negative control siRNA (AllStars, Qiagen) were used at a final concentration of 5 nM and mixed with 1.3 µl of Lipofectamine RNAiMAX (Thermo Fisher Scientific). The transfection mix was pre-spotted onto 24-well plates for 30 min at RT prior the addition of 7.0x10^4^ A549 cells per well. Infection experiments were carried out after 48 hours of transfection.

### Western Blot

Protein from cell lysates were prepared using the RIPA Buffer System (Santa Cruz Biotechnology) and FBS-free supernatants were concentrated using 3KDa centrifugal filter units (Thermo Fisher Scientific), following manufacturer’s instructions. BCA assay (Thermo Fisher Scientific) was used to determine protein concentration. Samples were normalized to the same concentration and separated under denaturing conditions on polyacrylamide gels (Bio-Rad). Proteins were transferred to polyvinylidene difluoride membranes (PVDF, Bio-Rad) and incubated for 1 h with 5% blotting grade blocker (Bio-Rad). Primary antibodies were incubated overnight at 4℃ and secondary HRP-conjugated antibodies were incubated for 1 h at room temperature. A list of all antibodies used is provided in supplemental Table S1. Super Signal West Pico (Thermo Fisher Scientific) was used for blot visualization by chemiluminescence using the ChemiDoc Touch Imaging System (Bio-Rad).

### RNA-immunoprecipitation and Co-immunoprecipitation

RNA immunoprecipitation (RIP) was performed as previously reported(9) with some modifications. At the indicated points, cells were scraped in RIP lysis buffer (200 mM KCl, 20 mM HEPES pH7.2, 2% N-dodecyl-β-D -maltoside, 1% Igepal, 100 U/mL Murine RNase inhibitor (NEB)). Lysates were kept on ice for 30 min and sonicated in an ultrasound bath cleaner (JP Selecta Ultrasons system, 40 kHz) for 15 sec.

Cleared lysates were centrifuged at 13,000 rpm for 15 min at 4℃ and supernatant recovered for protein quantitation. Protein was adjusted to 400 µg and incubated O/N at 4℃ with 5 µg of IP-antibody (Table S1) or rabbit IgG control (Abcam, Cambridge, UK) in a final volume of 500 µl of NT2 buffer (150 mM NaCl, 50 mM Tris-HCL pH 7.5, 1 mM MgCl_2_ and 0.05% Igepal). Dynabeads protein G (Invitrogen) were washed with NT2 buffer and blocked for 1 h with 1% BSA (Sigma-Aldrich, St. Louis, MO) and 50 µg of yeast RNA (Ambion). Blocked beads were added to the protein-antibody complex and kept at 4℃ for 2 h with constant rotation. Immunoprecipitation using the rabbit anti-C antibody were carried out in NT2-300 buffer, where the final concentration of NaCl was increased to 300 mM. Complexes were placed in a magnetic separation rack (NEB) to collect the unbound sample and then washed four times with NT2 buffer. After the last wash, complexes were resuspended in NT2 buffer and 10% of the sample was analyzed by western blot. The remaining sample was used for RNA extraction and quantification by RT-qPCR. For Co-IP experiments, cell lysates were prepared as described before with incubation and washes done with NT2-300 buffer. When indicated, IP samples were treated with RNase A/T1 (Ambion). In this case, IP samples were resuspended in 400 ul of RNase buffer (10 mM Tris-HCL pH 7.5, 300 mM NaCl and 5 mM EDTA). Half of the sample was left untreated and the other half treated with RNase A (2 µg/µl) and RNase T1 (5 U/µl); then samples were incubated for 1 h at 37℃. Samples were washed 3X with NT2-300 buffer and resuspended in 2% SDS for removal of dynabeads and consequent immunoblot analysis.

### Immunofluorescence and *in situ* proximity ligation assay (PLA)

Huh7 cells were grown on 12-mm coverslips to ∼50% confluency. Cells were infected with DENV-2 at the indicated MOI and time. At the end of an experiment, cells were fixed with 4% paraformaldehyde (PFA) and permeabilized with 0.1% SDS (Merck Millipore, Burlington, MA). Primary antibodies were incubated for 1 h at RT a humid chamber. Upon extensive washing, secondary antibody was added and incubation was continued for 1 h at RT in the dark. All antibodies were diluted in 0.5% milk powder (Bio-Rad) prepared in PBS. Coverslips were mounted with ProLong Gold Antifade mountant (Thermo Fisher Scientific), which contains DAPI. For PLA assays, cells were grown and fixed as described above and staining was conducted following the Duolink® PLA fluorescence protocol from Sigma-Aldrich. The indicated antibody and the PLA® probes anti-mouse minus and anti-rabbit plus were used. All samples were visualized by confocal microscopy (Leica TCS SP8 STED 3X laser scanning microscope equipped with a HC PL APO CS2 100x/1.40 oil objective). Laser intensity and detector gain were adjusted at the beginning of and remain unchanged for the duration of an experimental session. Scale bars in all images are 10 μm. All images were analyzed with ImageJ. The plug-in coloc2 was used for colocalization analysis and an in-house macro was designed to count the number of dsRNA foci and PLA signals per cell. At least 30 cells per experimental condition were analyzed in at least three independent experiments.

### Proteinase K (PK) protection assay

This assay was performed as indicated by Roder and Horner (32). Briefly, DENV-infected cells where harvested on ice-cold PK buffer (50 mM Tris, 10 mM CaCl2, 1 mM DTT) and permeabilized by five cycles of freeze thawing as described above. Protein samples were adjusted to the same concentration and divided into three treatments consistent of: non-treated negative control, triton X-100 and PK-treated positive control and PK-treated experimental sample. Cells were incubated with 10% triton X-100 (Sigma-Aldrich) for 5 min on ice and PK (Thermo Fisher Scientific) was used at a final concentration of 50 µg/ml for 30 min at RT. All tubes were treated with PMSF (Santa Cruz Biotechnology, Dallas, TX) at a final concentration of 1 mM for 10 min and processed for immunoblotting. The ratio of protein protected from PK-degradation was obtained by dividing the densitometry values of the PK-treated samples by that of the non-treated control and the levels of E protein degradation were calculated by dividing the densitometry value of the 70 KDa band by the densitometry value of the entire lane and subtracting this value from 100%.

### Transmission electron microscopy

Huh-7 wild type and YBX-1 KO cells were seeded into T75 flasks in regular growth media (DMEM, 10% FBS, 1% P/S). The following day cells were inoculated with DENV-2 NGC (MOI 1) at 37°C for two hours. After virus absorption cells were rinsed one time with PBS and fresh media (DMEM, 5% FBS, 1% P/S, 0.01 M Hepes) was added to the cells. 48 hours post infection media was removed, cells were rinsed with PBS and 2 ml of 0.25% trypsin-EDTA was added. After cells detached from the flask 8 ml of regular growth media was added and collected in 15 ml tubes. Cells were pelleted at 1000 rpm for 5 minutes. Media was removed and cells were rinsed 2X in room temperature PBS and fixed fixed for 30 mis in a mixture of 2.5% formaldehyde prepared from paraformaldehyde power, and 0.1% glutaraldehyde in in 0.05M cacodylate buffer pH 7.3 to which 0.01% picric acid and 0.03% CaCl_2_ were added. Cells were stored at 4°C until processed for TEM imaging. For ultrastructural analysis in (ultra)thin sections infected the monolayers were washed in 0.1M cacodylate buffer, cells were scraped off pelleted and processed further as a pellet. The pellets were post-fixed in 1% OsO_4_ in 0.1M cacodylate buffer pH 7.3 for 1 hr, washed with distilled water and *en bloc* stained with 2% acqueous uranyl acetate for 20 min at 60°C. The pellets were dehydrated in ascending concentrations of ethanol, processed through propylene oxide and embedded in Poly/Bed 812 (Polysciences, Warrington, PA). Ultrathin sections were cut on Leica EM UC7 ultramicrotome (Leica Microsystems, Buffalo Grove, IL), stained with lead citrate and examined in a JEM-1400 (JEOL USA, Peabody, MA) transmission electron microscope at 80 kV. Digital images were acquired with a bottom-mounted CCD camera Orius SC200 1 (Gatan, Pleasanton, CA).

## Statistical analysis

All data were analyzed with GraphPad Prism software. Differences between each of the YBX1 KO clones (KO1 and KO2) and WT cells were determined by non-parametric Mann-Whitney test. A p value of ≤0.05 was considered significant, with *p≤0.05, **p≤0.01 and ***p≤0.001. Data related to the analysis of confocal microscopy are shown as median + interquartile range (IQR) and all other data are presented as mean ± standard error of the mean (SEM).

